# Harnessing diversity and antagonism within the pig skin microbiota to identify novel mediators of colonization resistance to methicillin-resistant *Staphylococcus aureus*

**DOI:** 10.1101/2022.08.05.502505

**Authors:** Laurice Flowers, Monica Wei, Simon A. B. Knight, Qi Zheng, Aayushi Uberoi, Amy Campbell, Ting-Chun Jamie Pan, Jasmine Walsh, Erin Schroeder, Emily W. Chu, Charles W. Bradley, Raimon Duran-Struuck, Elizabeth A. Grice

## Abstract

The microbiota mediates multiple aspects of skin barrier function, including colonization resistance to pathogens such as *Staphylococcus aureus.* The endogenous skin microbiota limits *S. aureus* colonization via competition and direct inhibition. Novel mechanisms of colonization resistance are promising therapeutic targets for drug-resistant infections, such as those caused by methicillin-resistant *S. aureus* (MRSA). Here, we developed and characterized a swine model of topical microbiome perturbation and MRSA colonization. As in other model systems, topical antimicrobial treatment had little discernable effect on community diversity though the overall microbial load was sensitive to any type of intervention, including swabbing. In parallel, we established a porcine skin culture collection and screened 7,700 isolates for MRSA inhibition. Inhibitory isolates were represented across all major phyla of the pig skin microbiota and did not have a strong preference for inhibiting closely related species, suggesting that relatedness is not a condition of antagonism. Using genomic and phenotypic criteria, we curated 3 isolates to investigate whether prophylactic colonization would inhibit MRSA colonization in vivo. The 3-member consortium together, but not individually, provided protection against MRSA colonization, suggesting cooperation and/or synergy among the strains. These findings reveal the porcine skin as an underexplored reservoir of skin commensal species with the potential to prevent MRSA colonization and infection.

## INTRODUCTION

*Staphylococcus aureus* is a major cutaneous pathogen that also commonly asymptomatically colonizes nares and skin of healthy adults^1^. Colonization with *S. aureus* is a predisposing risk factor for infection and associated pathology^2,3^ and can precede a variety of different infections, ranging from local benign dermatopathologies (impetigo, folliculitis) to life-threatening abscesses, cellulitis, pneumonia, bone infection, and sepsis^4^. Of particular concern, methicillin-resistant *S. aureus* (MRSA) infections have increased markedly over the past 20 years and are associated with significant excess morbidity, mortality, and cost^5^. While *S. aureus* can colonize nares and skin asymptomatically, the factors that promote this pathogen’s shift from colonization to infection remain poorly understood.

*S. aureus* has also emerged as an important livestock pathogen, and causes infections in economically important animals such as sheep, goats, pigs, cows, rabbits, and chickens^6^. Livestock herds may act as a reservoir of newly virulent strains that cause infection in humans^7^. Pigs in particular have been implicated as a reservoir of MRSA and an incubator for antimicrobial resistance, due in part to the common practice of feeding subtherapeutic antibiotics to promote growth^8–10^. *S. aureus* strains that are pathogenic in livestock, specifically sequence type 398 (ST398; LA-MRSA), most likely originated in humans and later acquired antibiotic resistance cassettes after jumping to livestock hosts^11,12^. As antimicrobial resistance continues to deplete our therapeutic toolbox, there is an urgent need for novel treatment strategies, including those that are adaptable for human and livestock applications.

One factor that limits pathogen access and invasion is the skin’s endogenous microbiota, through a mechanism known as colonization resistance^13^. Previous work has shown that disruption of the skin microbiota with broad spectrum antimicrobials promotes colonization by *S. aureus*^14^. Competitive interactions that limit *S. aureus* skin colonization and pathogenicity are common among the coagulase-negative staphylococci (CoNS) species. For example, *S. hominis* and *S. lugdunensis* produce novel antibiotics that kill *S. aureus*^15,16^. Interference with quorum sensing is another mechanism of competitive exclusion, and is used by the CoNS species *S. caprae* to limit bacterial growth and fitness of *S. aureus*^17^. Additionally, skin commensals can stimulate and cooperate with host immune responses, thus playing an indirect role in excluding pathogens like *S. aureus.* Community-wide, these competitive interactions may in part explain how the commensal microbiota can exclude or counteract pathogens such as *S. aureus.*

Previous studies to characterize competitive interactions on skin have largely focused on CoNS, which are prominent members of the skin commensal microbiota as established by culture-independent and - dependent studies. The bias towards CoNS species could reflect their adaptation and competitive advantage on skin. It has also been postulated that competition is more common among closely related species^18^. However, this tendency could also reflect cultivation bias since CoNS are readily cultured from skin under the same conditions as *S. aureus* and MRSA. Recent studies have demonstrated that non-CoNS members of the skin microbiota also produce antibiotics, including *Cutibacterium acnes^19^* The extent to which skin commensals produce inhibitory molecules as means of competition is unknown, as is the link between phylogenetic relatedness and inhibitory capacity.

Model systems to isolate and study the skin microbiota often rely upon murine models. Fundamental differences in murine skin compared to human skin include morphological and histological differences such as thickness, the density of pilosebaceous units and other appendages such as sweat glands, and even distinct pathways and cell types involved in repair processes^20–22^. Furthermore, many *S. aureus* virulence factors are attenuated in mice.^23^ On the other hand, porcine skin displays similar morphology and histology to human skin and has similar density of appendages, and may represent a more suitable host for modeling skin microbial communities and MRSA colonization. *S. aureus* is also a natural pathogen of pigs, likely owing to similarities to human skin and nares. Because microbially-produced antibiotics are often specific to the range of species and environmental conditions likely to be encountered^24^, the pig skin microbiota may also harbor novel bacterial determinants and mechanisms of microbial antagonism. Inhibitory mechanisms applicable to livestock populations would have important implications for the veterinary control of *S. aureus,* in addition to the implications for human health and disease.

Here, we developed and characterized a pig skin model system to study microbial community perturbations and colonization resistance to MRSA. We first characterized the microbial diversity of porcine skin using cultures and 16S ribosomal RNA (rRNA) gene sequencing. We tested the resilience of the skin microbiota to pulse disturbances of various topical antimicrobial drugs, and then examined community resistance to MRSA colonization. This experimental design together with a high throughput in vitro inhibition screen led to the identification of 37 unique species that inhibited MRSA. We investigated the relationship between phylogenetic relatedness and antagonism to address whether closely related (e.g. *Staphylococcus)* species were more likely to display inhibition. Finally we colonized murine skin with 3 candidate inhibitors to test whether prophylactic colonization with pig commensal isolates also inhibited MRSA in vivo.

## RESULTS

### Bacterial composition of the dorsal skin microbiota of swine

We first characterized the baseline, unperturbed skin microbiota of Yucatan pig dorsal skin by analyzing 16S ribosomal RNA gene sequences. We demarcated 10 distinct squares on the dorsum each measuring ~40 cm^2^ (**Supplemental Figure S1**) and collected swabs from each square to sample the baseline skin microbiota. We observed at the phylum level similarities with the human bacterial skin microbiota, including a predominance of Firmicutes in the community (**Figure 1A**). At the genus level the most abundant taxa were *Streptococcus, Prevotella,* and *Lactobacillus* (**Figure 1B**). Genera that predominate on human skin were also found on porcine skin, albeit at lower relative abundances, including *Staphylococcus* and *Corynebacterium.* The effect of cohousing of the pigs was also apparent, as our studies were conducted over 4 experiments of 2 pigs each and batch effects are visible across paired individuals.

**Figure 1:**
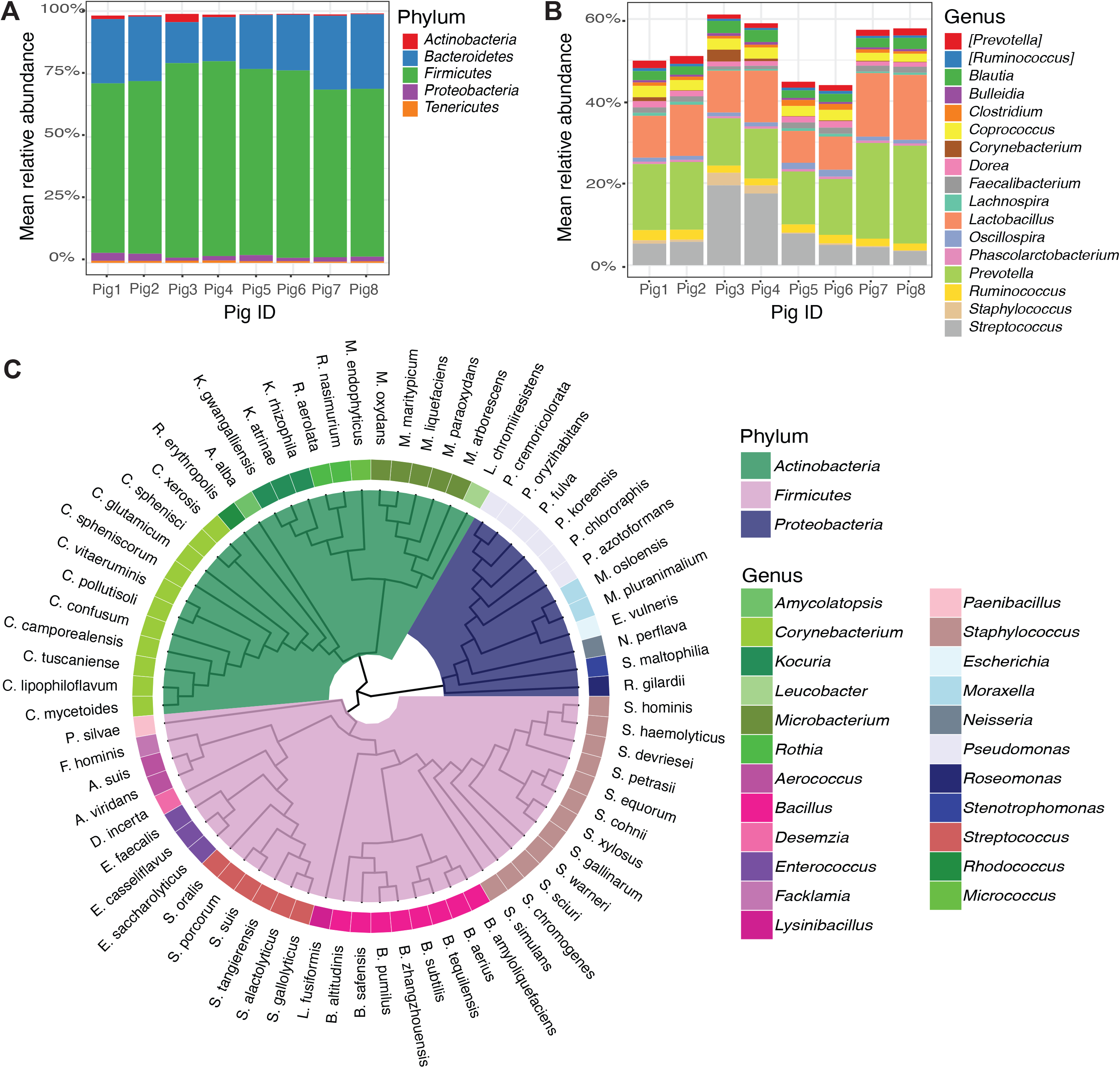
The culture-independent and -dependent microbiota of porcine skin. Mean relative abundance of top A) phyla and B) genera present in dorsal pig skin microbiota by 16S rRNA gene sequencing. Each bar represents an individual pig. C) Cladogram of bacterial isolates cultured from pig skin. Phylogeny was constructed from representative 16S rRNA gene sequences curated from the Silva database.

In parallel with culture-independent 16S rRNA amplicon sequencing, we characterized the culturable microbiota by plating skin swabs on blood agar under aerobic conditions. We identified 84 unique species or presumptive species using MALDI-TOF and full-length sequencing of the 16S rRNA gene for identification (**Supplemental Table S1**). Mirroring the culture-independent data, the top phylum recovered was Firmicutes, but we also cultured a substantial proportion of isolates that were identified as Actinobacteria and Proteobacteria (**Figure 1C**). Twelve species of coagulase-negative staphylococci were isolated, including *S. hominis, S. haemolyticus, S. warneri, S. cohnii,* and *S. simulans,* species that are also known to inhabit human skin. Eleven species of *Corynebacterium* were isolated including the food-industry staple *C. glutamicum,* the zoonotic pathogen *C. xerosis,* and human skin associated species such as *C. lipophiloflavum* and *C. mycetoides.* Overall, these results highlight some similarities between porcine and human skin microbiota, but also suggest a distinct cutaneous ecosystem.

### Community perturbation facilitates MRSA colonization on porcine skin

We previously observed in hairless (SKH-1 elite) mice that topical application of ethanol promoted *S. aureus* colonization^14^. We performed similar experiments using the swine model to test the effect of 8 topical perturbations on skin microbiota diversity and MRSA colonization. After collecting baseline microbiota specimens for characterization as described above, the 10 demarcated skin patches were treated twice daily for two days with topical interventions consisting of antiseptics (ethanol, betadine), antibiotics (mupirocin, triple antibiotic ointment), an anti-inflammatory (hydrocortisone), and an anti-fungal (clotrimazole) (**Figure 2A**). All ointments were prepared in a polyethylene glycol (PEG) vehicle, thus a PEG control was included in addition to two untreated controls. Swabs were collected for both culture-dependent and -independent profiling, twice daily.

**Figure 2:**
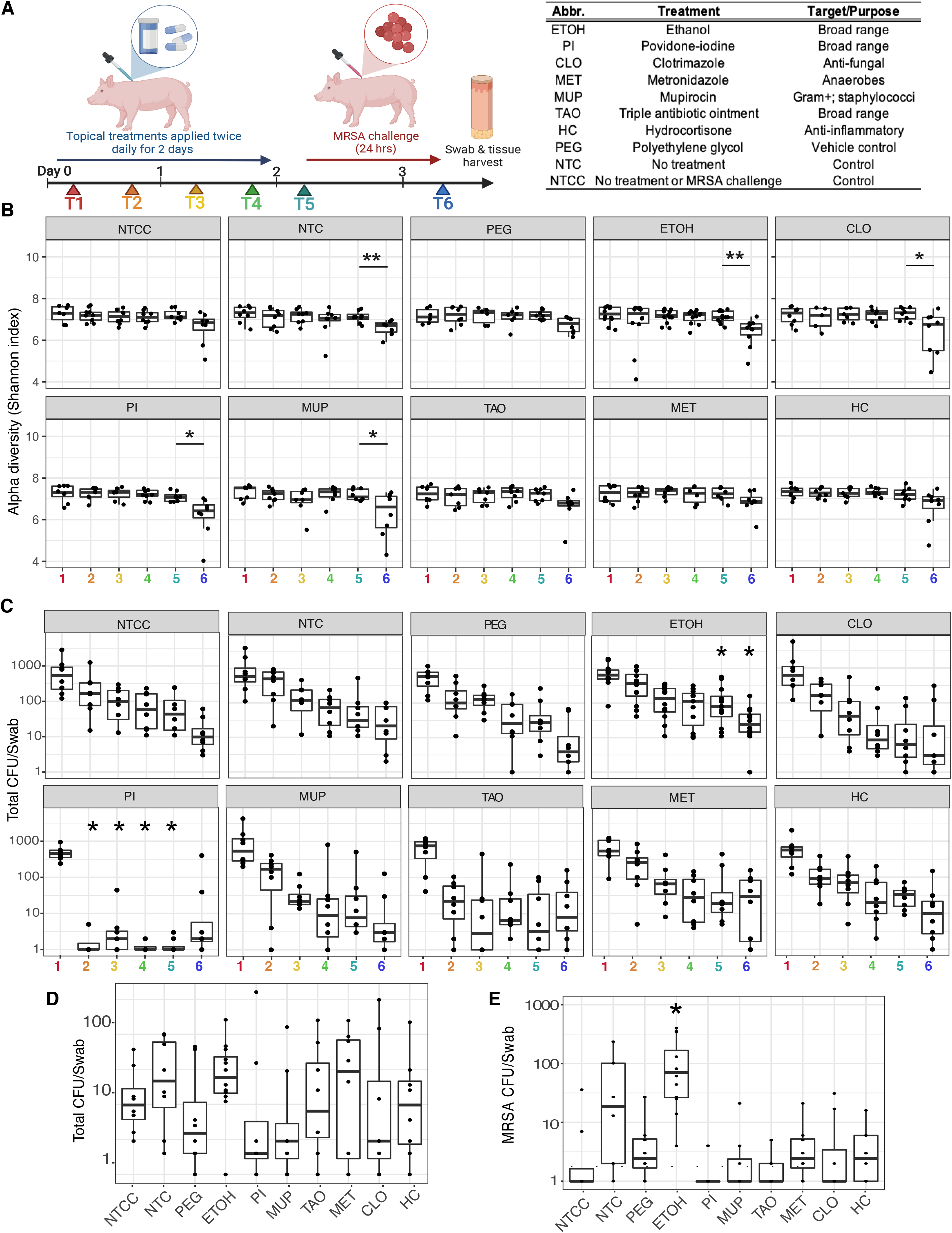
Skin microbiota dynamics during antimicrobial treatment and MRSA challenge. A) Experimental design for topical treatment and MRSA colonization. The accompanying table lists the different treatments and controls topically applied to skin, twice daily for two days. MRSA was then applied for 24 hours and tissue CFU was counted. T1-T6 (colored in rainbow) represent the timepoints where swabs were collected for 16S rRNA gene sequencing and culture quantification. B) Shannon diversity index (y-axes) represented over time, T1-T6 (x-axes). Each panel represents a different treatment, as indicated in the grey heading. Statistical testing was performed between T1 and all other timepoints for each treatment group individually using a t test; p values * < 0.05, **0.01. C) Total CFU recovered (y-axes) over time, T1-T6 (x-axes). Each panel represents a different treatment, as indicated in the grey heading. Statistical testing was performed between T1 and all other timepoints for each treatment group individually using a paired t-test with Bonferroni correction; * indicates p<0.05. D) Total CFU recovered (y-axis) at final T6 timepoint (treatment groups compared to NTC, Wilcoxon test with Bonferroni correction); E) CFU of MRSA (y-axis) recovered at T6 timepoint following the 24 hour challenge for each treatment (x-axis) (treatment groups compared to NTC control, Wilcoxon test with Bonferroni correction). A pseudocount of +1 was added to all CFU counts to accommodate the log scale. * indicates p<0.05.

We first examined how treatment altered the diversity of the skin microbiota, and found that the topical pulse disturbances had little discernable effect, as measured by the Shannon diversity index (**Figure 2B**). However, quantification of cultured CFU indicated a reduction of bacterial load on skin (**Figure 2C**). By quantifying culturable bacteria, we found in all cases including controls, there was a significant depletion of bacterial load. The most profound reduction was elicited by povidone-iodine, a broad range antiseptic that persists on the skin surface. The act of swabbing alone to collect endpoint data, as indicated by controls from untreated skin, was sufficient to reduce skin bacterial load as was treatment with the PEG vehicle, albeit not to the same extent as broad spectrum antimicrobials (**Figure 2C**). This suggests that mechanical disturbance by swabbing alone impacts the microbiome in ways that are not necessarily discernable by culture-independent methods.

To functionally characterize the skin microbiome configurations that result from topical pulse disturbance, each skin patch was challenged with 1×10^8^ CFU MRSA (USA300 Rosenbach strain) at timepoint 5 (T5) (**Figure 2A**). CFUs were quantified 24 hours later and microbial communities were profiled by 16S rRNA gene sequencing. MRSA challenge resulted in a significant decrease in community diversity when the skin was pre-treated with ethanol, povidone-iodine, hydrocortisone, as well as the vehicle PEG, as estimated by the Shannon diversity index (**Figure 2B**). The depletion in alpha diversity was consistent with an increase in the relative abundance of *Staphylococcus* by 16S rRNA amplicon sequencing at time point 6 (**Supplemental Figure S2**). Quantification of total CFU (**Figure 2D**) and MRSA CFU (**Figure 2E**) at the final time point indicated that ethanol treatment significantly enhanced MRSA colonization.

### Identification of phylogenetically diverse pig commensal isolates with anti-MRSA inhibitory activity

A total of 7,700 cultured skin isolates were spotted on a lawn of MRSA grown on blood agar to screen for inhibitory activity (**Figure 3A**). Of these isolates, 363 exhibited clearing, or a zone of inhibition (ZOI) surrounding the spot on the MRSA lawn. Identification of this group of isolates by MALDI-TOF mass spectrometry or sequencing of the 16S rRNA gene, indicated that it was comprised of 37 unique species (**Figure 3B**; **Supplemental Table S1**). Fifty-four isolates were not identified, and 58 were assigned at the genus level. *Aerococcus viridans* was the most common MRSA inhibiting isolate (40 isolates of the 363) followed by *Staphylococcus warneri* (26 isolates) and *Bacillus pumilus* (19 isolates). Six different species of *Staphylococcus* were identified all of which were coagulase-negative. Twenty-five of the isolates identified at the species level were represented by a single isolate, including a relatively undescribed bacterium *Desemzia incerta.* Not all the isolates were bacteria: single isolates of *Trichosporon asahii and Cryptococcus magnus* as well as three isolates of *Candida guilliermondii* were captured. The isolates of *C. magnus* and *C. guilliermondii* were from the same pig.

**Figure 3:**
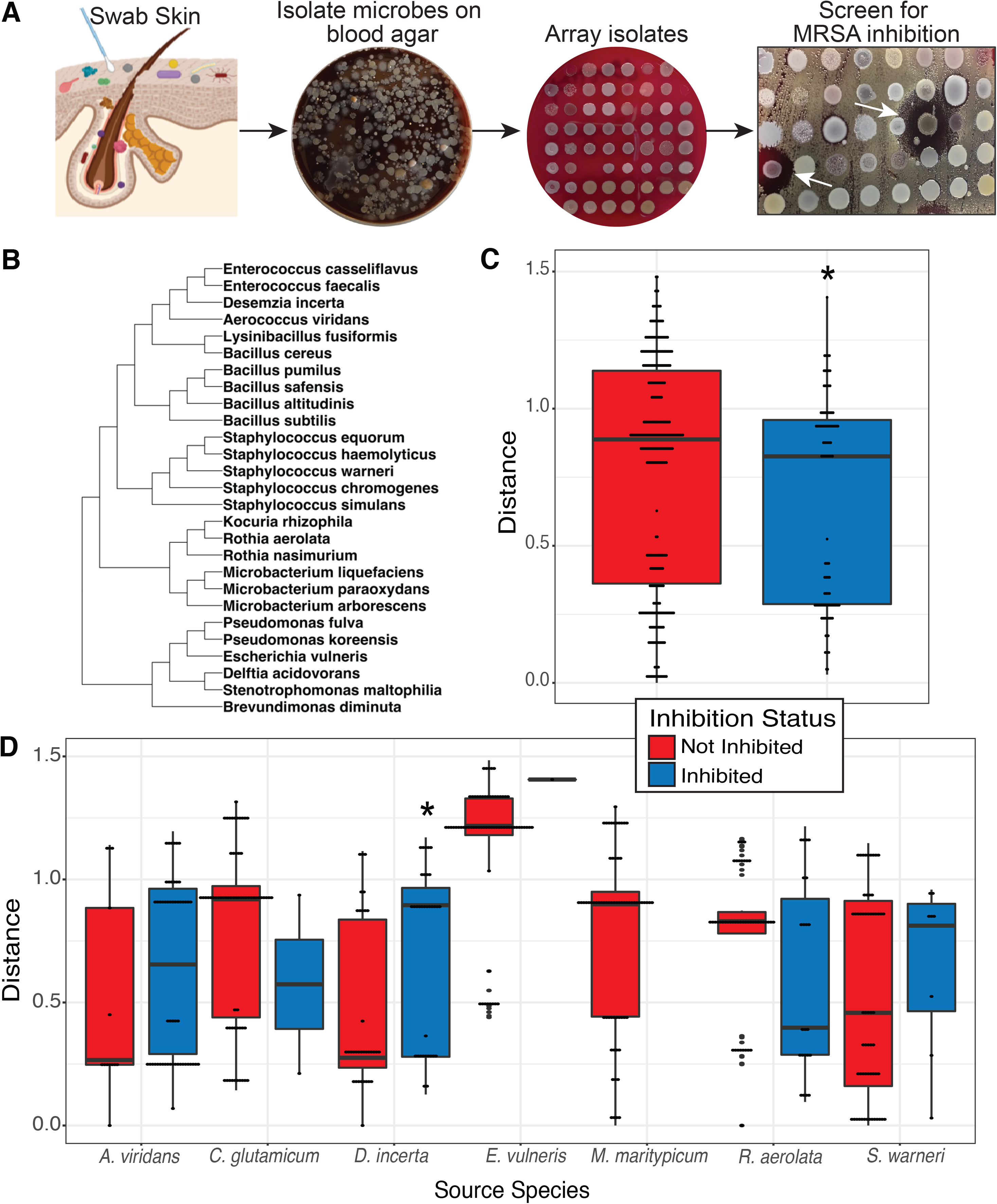
High throughput screening of pig skin commensal library identifies novel inhibitors of MRSA. A) Strategy for isolating and screening pig skin isolates for MRSA inhibitory activity; B. Cladogram of bacterial inhibitors of MRSA isolated from porcine skin; Phylogeny was constructed from representative 16S rRNA sequences from the Silva database; C. Patristic distance between bacterial isolate pairs that showed an inhibitory interaction (red) vs. a non-inhibitory interaction (blue) on agar diffusion assay (Wilcoxon test). D. Patristic distance between bacterial isolate pairs that showed an inhibitory interaction vs. non-inhibitory interaction, separated by “Source” species (inhibited vs. non-inhibited for each group; Wilcoxon test).

### Phylogenetic distance is not strongly correlated with antagonism

Studies of environmental microbial communities have demonstrated a correlation between phylogenetic distance and microbial antagonism^18^. In line with this finding, previous studies of competitive microbial interactions on skin have largely identified CoNS as inhibitors of *S. aureus.* Surprisingly, only 6/37 (16.2%) of MRSA inhibitors that we isolated from porcine skin were *Staphylococcal* species, suggesting that phylogenetically distant inhibitors of *S. aureus* may be common in some microbial communities. To test whether bacteria from porcine skin showed a preference for inhibiting phylogenetically similar species, we selected seven isolates (“Source” species) for further characterization of inhibition patterns. Five isolates – *D. incerta, A. viridans, R. aerolata, E. vulneris,* and *S. warneri* – were inhibitors of MRSA, while two isolates – *C. glutamicum* and *M. maritypicum* were not inhibitors of MRSA. Using the agar diffusion assay described above, we performed pairwise inhibition testing of these seven isolates against the remaining 68 isolates (“Target” species) in the porcine skin commensal collection. We tested for inhibition on two growth medias (BHI-T agar and blood agar) and two temperature conditions (37°C and room temperature, ~22-23°C). An interaction was considered inhibitory if the target species was inhibited in at least one growth condition. A total of 1553 assays were performed across all growth conditions which were consolidated into 469 pairwise interactions.

For each pairwise interaction, we estimated phylogenetic distance based on a multiple sequence alignment of representative, publicly available 16S rRNA gene sequences. Across all interactions, we found smaller mean cophenetic distance between inhibitory pairs compared to non-inhibitory pairs (**Figure 3C**; p =0.016). This finding is consistent with other studies demonstrating a higher probability of bacterial antagonism between phylogenetically related species. However, when interactions were separated by source species, this trend was not reproduced. Six out of 7 species showed no statistically significant relationship between phylogenetic distance and inhibitory interaction. *D. incerta* showed higher mean phylogenetic distance between inhibited species compared to non-inhibited species (**Figure 3D**; p=0.029). Thus, while a relationship between phylogenetic distance and bacterial inhibition seems to exist at the broader community level, individual bacterial species exhibited a diverse range of behaviors.

### A consortium of inhibitory bacteria provides protection against MRSA colonization in vivo

We further focused on three inhibitory species in particular, *Aerococcus viridans, Desemzia incerta,* and *Rothia aerolata,* (**Figure 4A**). These were selected because they are phylogenetically diverse, inhibited MRSA under distinct conditions, and were predicted to use different mechanisms of inhibition. Genomic sequencing of the 3 isolates enabled prediction of biosynthetic gene clusters (BGCs), comprising the genes needed to produce, process, and export small molecules and peptides that mediate interactions between species. This analysis indicated non-overlapping sets of BCGs between the 3 isolates (**Figure 4B**). R*. aerolata* contained BGCs predicted to encode a lanthipeptide and a betalactone, *D. incerta* BGCs included a type 3 polyketide synthase (T3PKS) and a terpene, and *A. viridans* BGCs included a bacteriocin and a terpene. *A. viridans* and *D. incerta* additionally inhibited *Pseudomonas aeruginosa,* and *D. incerta* also inhibited *Streptococcus pyogenes* (**Supplemental Figure S3**). Thus, these 3 isolates are likely to employ non-overlapping mechanisms of inhibition.

**Figure 4:**
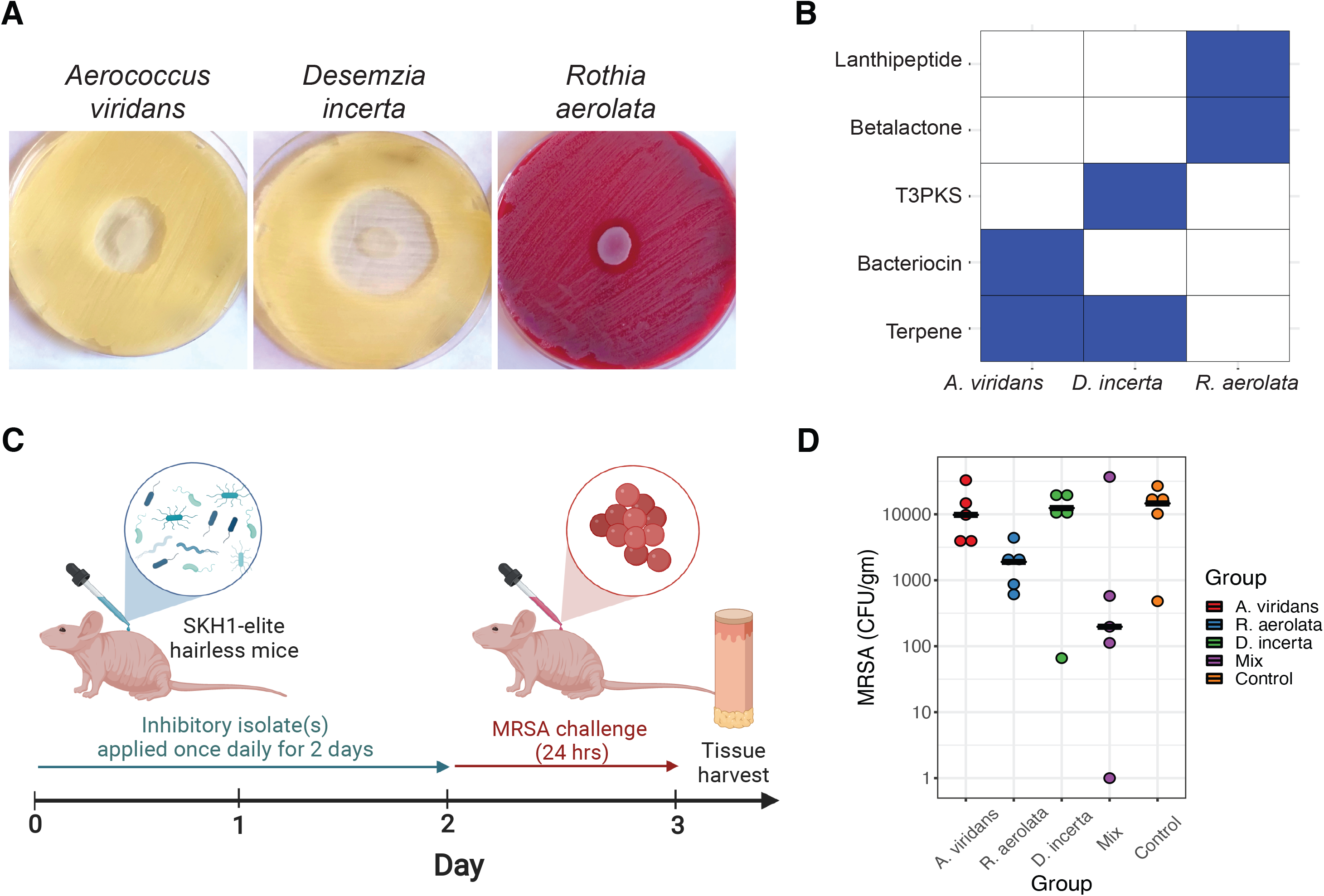
A defined consortium of pig skin isolates reduces MRSA colonization on murine skin. A) Photographs of MRSA inhibition for each of the three selected isolates. USA300 strain MRSA*, Aerococcus viridans, Desemzia incerta,* and *Rothia aerolata* were grown overnight in BHI+0.8%Tween80 media. MRSA was diluted to OD_600_ of 0.1 and spread onto BHI-T agar (for *A. viridans* and *D. incerta)* or TSA + 5% blood agar (for *R. aerolata).* Pig skin isolates equivalent to an OD_600_ of 2.0 were spotted onto the MRSA lawn. B) Biosynthetic gene cluster profiles based on genomic data for each isolate. C) Schematic of experimental design for murine colonization experiments. D) CFU of MRSA recovered (y-axis) when skin is pre-colonized with the indicated isolate, a mix of all 3, or control. N=5 mice per group. Bars indicate the median for each treatment group. Statistical testing performed comparing treatment group to Control using Wilcoxon test with Bonferroni correction. A pseudocount of +1 was added to all CFUs to accommodate the log scale.

While the three porcine skin isolates we selected inhibit MRSA in vitro, it was uncertain whether these isolates would provide the same protection in vivo. Furthermore, since the mechanisms of inhibition were likely different, we hypothesized that using the 3 isolates in a mix (3-Mix) would provide more protection than monocolonization with a single inhibitory isolate. To test this, we returned to a hairless mouse model we previously used in studies of skin colonization resistance^14^. SKH-1 elite mice were colonized prophylactically with each isolate or the 3-Mix daily for 2 days (**Figure 4C**). On the 3^rd^ day, skin was challenged with MRSA (1×10^8^). After 24 hours, skin was collected and processed to quantify MRSA CFU and to calculate colonization efficiency. We performed a pilot study with 4.5×10^7^ inoculum of each pig isolate and found that individually, no single isolate was consistently effective at disrupting MRSA colonization (**Supplemental Figure S4**). However, the 3-Mix blocked MRSA colonization in 3 of the 4 mice, to a median 733 CFU per gram of skin, compared to a median 1.06 x 10^4^ CFU per gram skin in the control. Because there was a large amount of variability between mice, we repeated the experiment with a higher inoculum of the pig commensal isolates (1×10^8^ each isolate). Similar to the pilot experiment, 4 of the 5 mice treated with the 3-Mix had ~ 2 log decrease in MRSA colonization compared to the controls (median=1.96 x10^2^ and 1.46 x 10^4^ CFU, respectively) (**Figure 4D**). While the differences we observed were not statistically significant, the consistency of the findings over independent experiments suggests that the 3-Mix is more effective than each component individually in providing protection against MRSA colonization in this model.

## DISCUSSION

Here, we developed and employed a porcine skin microbiome model to identify novel bacterial determinants of colonization resistance to the skin and soft tissue pathogen MRSA. Combining culture-dependent and -independent approaches, we find that the pig skin microbiota has some similarities to human skin communities, including a phylogenetic preference for Actinobacteria and Firmicutes. We cultured and screened 7,700 pig skin isolates and identified 363 with inhibitory activity against MRSA. The majority of these inhibitory isolates were non-Staphylococcal species and could inhibit other bacteria across a broad phylogenetic range. Combining phenotypic and genomic analysis, we selected 3 distinct isolates to examine their capacity to inhibit MRSA in vivo. We found that the best protection was provided by a consortium of the 3 inhibitory isolates and that monocolonization with individual inhibitors was not as effective.

Our rationale for pursuing novel bacterial determinants of colonization resistance in the pig model is three-fold: 1) pig skin resembles human skin more than other commonly used model organisms; 2) livestock, including swine, can be infected and act as reservoirs for *S. aureus* and MRSA; 3) the understudied ecosystem of the pig microbiome may contain novel mediators of colonization resistance that could be used to combat drug-resistant infections like MRSA. The pig model also has some disadvantages, such as the cost and labor associated with husbandry, and the unavailability of tools and reagents (e.g. antibodies) to study host-related factors. In this case, the utility of the pig model was most apparent in the initial screen to identify novel isolates that inhibit *S. aureus.*

Previous to this work, the majority of skin microbial strains that have been investigated for *S. aureus* inhibition were other staphylococcal species^13^. We speculated that this might reflect the ease of isolation and dominance in the human skin microbiota, rather than a phylogenetic bias of closely related species inhibiting each other. In line with previous studies from soil microbes, we found that the porcine skin isolates we examined were more closely related to species they inhibited than to non-inhibited species. However, the effect size between groups was very small (0.11, approximately the distance between two species in the same genus). We found that bacterial species could inhibit other species across a broad phylogenetic range. We furthermore identified a number of *S. aureus* inhibitors that were not *Staphylococcus* species. These findings suggest that genetically dissimilar bacteria still show a high rate of inhibition and should not be excluded from a search for inhibitory species or antimicrobial products.

Even though the 3-Mix of pig inhibitory isolates provided the most promising results for prophylactic colonization against MRSA, there was still a large amount of variability between mice that precluded statistical significance. The transplant of exogenous species to the skin can be hampered by the endogenous microbiota, in this case of the hairless mice. Resource exclusion and/or antagonism by the microbiota, as well as host factors such as habitat filtering and immune mechanisms, are important factors that can influence the engraftment and function of a microbiota transplant^25^. Thus, future studies will need to address these factors through an ecological framework, potentially by using germ free and gnotobiotic mice, as well as investigating the impact of the inhibitory isolates on host immune response.

Previous studies have explored the potential of the human commensal microbiota, and strains within, to combat *S. aureus* colonization and infection^15–17,26,27^. As antibiotic resistant bacteria present an ever-increasing threat to our diminishing supply of antimicrobial drugs, alternative modes of therapy, such as bacteriotherapy, may offer promising opportunities to combat colonization and prevent infections. Our work demonstrates that similar to colonization resistance to vancomycin-resistant *Enterococcus* in the gut ^28^, a consortium of strains was required to provide protection to pathogen colonization. Other factors, such as the endogenous microbiota and the host may play a role in how transplanted communities function. Furthermore, leveraging the novel microbial-microbial interactions within underexplored microbial ecosystems that encounter similar threats (i.e. MRSA) may lead to novel antibiotic targets.

## METHODS

### Housing and care of pigs

Eleven, three to three and half month-old female Yucatan pigs (Sinclair Bio Resources LLC), weighing between 12 to 17 Kg were housed in individual pens with unlimited access to food and water in a USDA approved ABSL2 facility. The pigs were acclimatized for three days, and then were socialized for handling, by being fed cake-frosting and dog biscuits whilst being confined with a pig-board twice daily for one week. Three pigs were used in pilot experiments and eight pigs were used for the main experiment. All experiments adhered to the regulations of the Animal Welfare Act and were approved by the University of Pennsylvania Institutional Animal Care and Use Committee (IACUC) IRB #806346

### Swab collection, application of topical ointments, and MRSA

The dorsal region of the pig was marked into ten 5 x 8 cm regions with permanent skin ink (**Supplemental Figure S1**). Four skin swabs from each region were taken, with each swab covering ¼ of the region area (10 cm^2^), rotating swab collected in the specific patch at each sampling. Swabbing was performed using a sterile foam tip applicator (Puritan), moistened in PBS (Corning), rubbed back and forth and crosswise with firm pressure for 10 seconds. The tip of one swab was broken off into a 2 ml tube (BioPur, Eppendorf) and snap frozen in dry-ice and stored at −80°C. The tip of the remaining 3 swabs were broken off into a 1.5 ml tube (BioPur, Eppendorf) containing 0.3 ml sterile PBS to isolate microbiota.

Drug ointments were prepared by dissolving each compound (**Figure 2A**) in 90% polyethylene glycol (PEG 3350) and applied with a sterile foam tip applicator to each marked region.

MRSA (ATCC #BAA-1717, Rosenbach strain of community-acquired USA300), grown at 37°C, 18 h, 250 rpm in tryptic soy broth (TSB, BD), and diluted to give 1×10^8^ CFU in 0.5 ml PEG, was applied to marked regions of the pig’s skin with a sterile foam tip applicator.

### Microbiome sequencing and analysis

#### DNA Extraction

Bacterial DNA was extracted from swabs and stored at −80°C, as described previously [Meisel et al 2016]. Swabs were incubated for one hour at 37°C with shaking in 300μL yeast cell lysis solution (from Lucigen MasterPure Yeast DNA Purification kit) and 10,000 units of ReadyLyse Lysozyme solution (Lucigen). Samples were subjected to bead beating for ten minutes at maximum speed on a vortex mixer with 0.5 mm glass beads (MoBio), followed by a 30-minute incubation at 65°C with shaking. Protein precipitation reagent (Lucigen) was added and samples were spun at maximum speed. The supernatant was removed, mixed with isopropanol and applied to a column from the PureLink Genomic DNA Mini Kit (Invitrogen). Instructions for the Invitrogen PureLink kit were followed exactly, and DNA was eluted in 50 μL elution buffer (Invitrogen). At each sampling event, swab control samples that never came into contact with the skin were collected, prepared and sequenced exactly as the experimental samples. No significant background contamination from either reagent and/or collection procedures was recovered.

#### Sequencing and analysis

Amplification of the 16S rRNA gene V1–V3 region was performed as described previously^29^. Sequencing was performed at the PennCHOP microbiome core on the Illumina MiSeq using 300 bp paired-end chemistry. The mock community control (MCC; obtained from BEI Resources, NIAID, NIH as part of the Human Microbiome Project: Genomic DNA from Microbial Mock Community B (Even, Low Concentration), v5.1L, for 16S rRNA Gene Sequencing, HM-782D) was sequenced in parallel. Sequencing of the V1-V3 region was performed using 300 bp paired-end chemistry. Sequences were preprocessed and quality filtered prior to analysis, including size filtering to 460-600 nucleotides. HmmUFOtu was used for sequence alignment and phylogeny-based OTU clustering as described previously^30^. Statistical analysis and visualization was performed using the phyloseq package^31^ in the R statistical computing environment. Amplicon sequences are publicly available in the NCBI SRA, BioProject Accession: PRJNA866157

### Porcine bacteria isolation

Tubes containing skin swabs in PBS were vortexed vigorously for 10 mins at room temperature (RT). Then 100 μl from each sample was spread onto two blood agar (BA) plates. The initial swab samples (T1, Fig 2A) were diluted 1:10 in PBS prior to plating. The BA plates were incubated at 37°C, 5% CO2 overnight, and RT for three days. Morphologically unique looking colonies from each plate were picked and transferred to a second BA plate, that was incubated at 37°C overnight and used as the source plate for the screen below.

### Screen for MRSA inhibiting isolates

Isolates from the blood agar source plates were inoculated into Brain Heart Infusion medium containing 0.8 % Tween 80 (BHI-T) in 96-well plates. After 24 h at 37°C, cultures were resuspended and diluted 1:10 into fresh BHI-T. Following overnight incubation at 37°C a 96-pin replicator was used to transfer 0.2 μl of each culture to blood agar (BA) plates on which 100 μl of an overnight culture of MRSA diluted to OD_600_ = 0.1 had been spread. Control plates without MRSA were also inoculated. The plates were incubated at 37°C for 24 hours, photographed, and colonies that inhibited the growth of MRSA, identified by a ring of no MRSA growth around the isolate, were selected.

### MALDI-TOF MS identification of MRSA inhibiting porcine isolates

Isolates that exhibited inhibition of growth of MRSA were streaked for single colonies on BA plates and incubated at 37°C overnight. Bacterial identification was performed using the MALDI Biotyper Microflex LT System (Bruker Daltonik GmbH) and the accompanying library, MBT BDAL 8468 MSP Library. According to the manufacturer’s instructions a genus and species level identification is accepted with a Maldi-TOF MS score of > 2.00 and a genus level identification is accepted at a score of 1.75-1.99 when backed by other ancillary microbiological identification methods. Formic acid was applied to all Gram-positive organisms in the run to breakdown the cell wall. Full tube extraction method was utilized for some sample identifications. All samples were run in duplicate.

### 16S rRNA gene Sanger sequencing

Colony PCR using GoTaq DNA polymerase (Promega) was performed on isolates not identified by MALDI-TOF MS. Primers (8F [AGAGTTTGATCCTGGCTCAG] and 1391R [GACGGGCGGTGTGTRCA]^32^ amplified the 16S rRNA gene, with an initial heating of 98°C 3 min followed by 35 cycles of 95°C 45 sec, 50 to 58°C 60 sec, 72°C 90 sec. PCR reactions were cleaned using Exo-CIP™ (NEB) and sequenced on an ABI 8730XL with BigDye Taq FS Terminator V3.1 (University of Pennsylvania, Penn Genomic Core). The DNA sequence was compared against the bacteria database of NCBI using the default settings of blastn.

### Construction of phylogenetic tree

Cultured bacterial isolates that could be definitively identified to the species level were included in the phylogenetic analysis. Representative 16S ribosomal RNA sequences for each species were curated from the Silva Living Tree Project (LTP) database. For species without a Living Tree Project entry, the longest high-quality sequence from the Reference Non-Redundant (refNR) dataset was used.

Multiple sequence alignment of representative 16S rRNA sequences was generated using MAFFT v. 7.505 using the L-INS-i setting. A maximum likelihood tree was constructed from the multiple sequence alignment with RAxML v. 8.2.12 using the best tree from 100 searches. The tree was midpoint rooted in FigTree v. 1.4.4 and visualized in RStudio. Pairwise cophenetic distances between species were calculated using the R ape package v 5.5.

### Bacteria-Bacteria Interaction Screen

Eight MRSA inhibiting isolates were tested against a collection of 88 pig isolates (37 of which could inhibit the growth of MRSA) in a growth inhibition spot diffusion assay. Single isolate colonies, which had been grown on BA plates for 72h, RT were inoculated into TSB and incubated at RT. After 72 h the TSB cultures were diluted to an OD_600_ = 0.1 in PBS and 60 μl of the cell suspensions were spread onto BA and BHI-T agar plates. The TSB cultures of the eight isolates to be spotted onto the lawns were diluted to OD_600_ = 1.0 in PBS. For *D. incerta,* which grows to a low cell density, it was sometimes necessary to pellet the cells (10,000 x g, 5 min, RT) and resuspend the cell pellet in a lower volume of PBS to achieve a cell suspension with an OD_600_ of 1.0. Five microliters of the cell suspensions were spotted onto the bacteria lawns and control plates. The plates were incubated at RT for 72 h and 37°C for 24 h, at which point they were photographed and zones of inhibition recorded.

Bacterial isolates that had not been identified to species level by either MALDI-TOF or 16S rRNA sequencing were excluded from the in silica analysis. For all growth conditions, each bacterial interaction was graded as inhibitory (1) or non-inhibitory (0) based on the presence of a visible zone of inhibition. Interactions across growth conditions were then consolidated such that a bacterial interaction was considered inhibitory if inhibition was observed in at least one growth condition. A phylogenetic tree of all isolates was constructed from representative 16S rRNA sequences as described above. Pairwise cophenetic distances between species in the tree were calculated using the R ape package v 5.5.

### Whole genome sequencing of bacterial genomes

Bacterial isolates from frozen stock were streaked onto blood agar plates. A single bacterial colony was inoculated into 3 mL Tryptic Soy Broth (TSB) media and incubated at 37C overnight without shaking. Genomic DNA was extracted using the ZymoResearch QuickDNA Fungal/Bacterial MicroPrep kit. Genomic DNA was sequenced by the PennCHOP Microbiome Core on the Illumina HiSeq2500.

### Genome assembly and annotation of biosynthetic gene clusters

Quality control of raw genomic reads was performed using FastQC v.0.11.8. The Nextera adapter sequence and 10 additional base pairs from each paired read were trimmed using TrimGalore v. 0.6.5. Quality control of trimmed reads was again performed using FastQC v.0.11.8. Genomes were assembled *de novo* with Unicycler v0.4.8 using standard settings. Whole genome sequences of selected commensal bacteria were mined for biosynthetic gene clusters with antiSMASH v 5.1.2 using standard settings.

### Murine colonization experiments

All mouse procedures were performed under protocols approved by the University of Pennsylvania IACUC. Seven week old female SKH1-elite mice were purchased from Charles River (#477) and allowed to acclimate for 1 week before experimentation began. Mice were given ad libitum access to food and water. Pig commensal isolates were grown in liquid TSB at RT for 24 hours at 300 rpm, then 37C for 24 hours at 300 rpm. On the following day, OD600 measurement was used to standardize inoculums, and pellets resuspended in TSB to acquire 2×10^9^ CFU/ml inoculums. Titers were validated by culture and OD600 measurements. Mice were monoassociated at the dorsum by pipetting on 50 ul of pig isolate inoculum then spread using a sterile swab. Applications of pig commensal suspensions was repeated 24 hours later. MRSA (ATCC #BAA-1717; USA300 Rosenbach strain) was grown in TSB overnight at 37C with 300 rpm shaking, with inoculum calculated and diluted similarly. A volume of 50 ul (of 2×10^9^ CFU/ml inoculum) was applied 24 hours post-association with the pig commensal isolate(s).

### Statistical Analysis and Data Visualization

All statistical analysis was performed using functions built into the R statistical environment (Rstudio version 1.4.1106). Data were visualized using ggplot2^33^ and ggtree.^34^ Normality of data was assessed by the Shapiro-Wilk test and visually by QQ-plot. T-test was used for groups that did not show evidence of nonnormality by the Shapiro-Wilk test (Fig. 2C). Wilcoxon test was used for data whose distribution departed significantly from normality (Fig 2D, 2E, 3C, 2D, 4D) based on the Shapiro-Wilk test. Bonferroni correction for multiple hypothesis testing was applied where applicable (Fig 2C, 2D, 2E, 3D, 4D). The distribution for 2 out of 10 treatment groups (TAO and PI) in Figure 2C departed significantly from normality based on Shapiro-Wilk test. We believe this occurred due to the high number of 0 values resulting in sparse data. The distribution of the remaining 8 out of 10 treatment groups did not show evidence of non-normality by the Shapiro-Wilk test. A parametric t-test was applied to all groups in Fig. 2C.

## ACKNOWLEDGEMENTS

We thank the veterinary staff, residents, and faculty of the University Lab Animal Resources (ULAR) at UPenn for their guidance and support in designing and optimizing the pig model, as well as their care for the animals; Penn Vet PADLS New Bolton Center Clinical Microbiology Lab for MALDI-TOF identifications (PI: Donna Kelly; Technical Staff: Denise Barnhart and Sean Loughrey); Joseph Horwinski, Julia Bugayev, and Emilio Rodriguez for technical assistance; current and former members of the Grice lab and the Department of Dermatology for critical discussion and review of the work. This work was funded by the following grants to EAG from the NIH, NIAMS (R01AR006663), NINR (R01NR015639), the Burroughs Wellcome Fund PATH Award, the University of Pennsylvania Linda Pechenik Montague Investigator Award, and the Dermatology Foundation Sun Pharma Research Award. This research was also supported by the Penn Skin Biology and Disease Resource-based Center (Penn SBDRC supported by NIH/NIAMS P30AR069589). LF was supported by the Penn Dermatology Research T32 Training Grant (NIH/NIAMS T32AR007465); MW was supported by the Bacterial Pathogenesis T32 Training Grant, (NIH/NIAID 5T32AI141393); AU is supported by the Prevent Cancer Foundation Awesome Games Done Quick fellowship; AC was supported by the UPenn Blavatnik Family Fellowship.

## REFERENCES

1. Olaniyi, R., Pozzi, C., Grimaldi, L. & Bagnoli, F. Staphylococcus aureus-Associated Skin and Soft Tissue Infections: Anatomical Localization, Epidemiology, Therapy and Potential Prophylaxis. Curr Top Microbiol Immunol 409, 199–227 (2017).

2. von Eiff, C., Becker, K., Machka, K., Stammer, H. & Peters, G. Nasal carriage as a source of Staphylococcus aureus bacteremia. Study Group. N Engl J Med 344, 11–16 (2001).

3. Wertheim, H. F. L. et al. Risk and outcome of nosocomial Staphylococcus aureus bacteraemia in nasal carriers versus non-carriers. Lancet 364, 703–705 (2004).

4. Tong, S. Y. C., Davis, J. S., Eichenberger, E., Holland, T. L. & Fowler, V. G. Staphylococcus aureus infections: epidemiology, pathophysiology, clinical manifestations, and management. Clin Microbiol Rev 28, 603–661 (2015).

5. Kourtis, A. P. et al. Vital Signs: Epidemiology and Recent Trends in Methicillin-Resistant and in Methicillin-Susceptible Staphylococcus aureus Bloodstream Infections - United States. MMWR Morb Mortal Wkly Rep 68, 214–219 (2019).

6. Fitzgerald, J. R. Livestock-associated Staphylococcus aureus: origin, evolution and public health threat. Trends Microbiol 20, 192–198 (2012).

7. Cuny, C., Wieler, L. H. & Witte, W. Livestock-Associated MRSA: The Impact on Humans. Antibiotics (Basel) 4, 521–543 (2015).

8. Ferguson, D. D., Smith, T. C., Hanson, B. M., Wardyn, S. E. & Donham, K. J. Detection of Airborne Methicillin-Resistant Staphylococcus aureus Inside and Downwind of a Swine Building, and in Animal Feed: Potential Occupational, Animal Health, and Environmental Implications. J Agromedicine 21, 149–153 (2016).

9. Sieber, R. N. et al. Drivers and Dynamics of Methicillin-Resistant Livestock-Associated Staphylococcus aureus CC398 in Pigs and Humans in Denmark. mBio 9, e02142–18 (2018).

10. Looft, T. et al. In-feed antibiotic effects on the swine intestinal microbiome. Proc Natl Acad Sci U S A 109, 1691–1696 (2012).

11. García-Álvarez, L. et al. Meticillin-resistant Staphylococcus aureus with a novel mecA homologue in human and bovine populations in the UK and Denmark: a descriptive study. Lancet Infect Dis 11, 595–603 (2011).

12. Price, L. B. et al. Staphylococcus aureus CC398: host adaptation and emergence of methicillin resistance in livestock. mBio 3, e00305–11 (2012).

13. Parlet, C. P., Brown, M. M. & Horswill, A. R. Commensal Staphylococci Influence Staphylococcus aureus Skin Colonization and Disease. Trends Microbiol 27, 497–507 (2019).

14. SanMiguel, A. J., Meisel, J. S., Horwinski, J., Zheng, Q. & Grice, E. A. Topical Antimicrobial Treatments Can Elicit Shifts to Resident Skin Bacterial Communities and Reduce Colonization by Staphylococcus aureus Competitors. Antimicrob Agents Chemother 61, e00774–17 (2017).

15. Zipperer, A. et al. Human commensals producing a novel antibiotic impair pathogen colonization. Nature 535, 511–516 (2016).

16. Nakatsuji, T. et al. Development of a human skin commensal microbe for bacteriotherapy of atopic dermatitis and use in a phase 1 randomized clinical trial. Nat Med 27, 700–709 (2021).

17. Paharik, A. E. et al. Coagulase-Negative Staphylococcal Strain Prevents Staphylococcus aureus Colonization and Skin Infection by Blocking Quorum Sensing. Cell Host Microbe 22, 746–756.e5 (2017).

18. Russel, J., Røder, H. L., Madsen, J. S., Burmølle, M. & Sørensen, S. J. Antagonism correlates with metabolic similarity in diverse bacteria. Proc Natl Acad Sci U S A 114, 10684–10688 (2017).

19. Claesen, J. et al. A Cutibacterium acnes antibiotic modulates human skin microbiota composition in hair follicles. Sci Transl Med 12, eaay5445 (2020).

20. Hamilton, D. W. et al. The pig as a model system for investigating the recruitment and contribution of myofibroblasts in skin healing. Wound Repair Regen 30, 45–63 (2022).

21. Sullivan, T. P., Eaglstein, W. H., Davis, S. C. & Mertz, P. The pig as a model for human wound healing. Wound Repair Regen 9, 66–76 (2001).

22. Duran-Struuck, R. et al. Miniature Swine as a Clinically Relevant Model of Graft-Versus-Host Disease. Comp Med 65, 429–443 (2015).

23. Trübe, P. et al. Bringing together what belongs together: Optimizing murine infection models by using mouse-adapted Staphylococcus aureus strains. Int J Med Microbiol 309, 26–38 (2019).

24. Hibbing, M. E., Fuqua, C., Parsek, M. R. & Peterson, S. B. Bacterial competition: surviving and thriving in the microbial jungle. Nat Rev Microbiol 8, 15–25 (2010).

25. Walter, J., Maldonado-Gómez, M. X. & Martínez, I. To engraft or not to engraft: an ecological framework for gut microbiome modulation with live microbes. Curr Opin Biotechnol 49, 129–139 (2018).

26. Brown, M. M. et al. Novel Peptide from Commensal Staphylococcus simulans Blocks Methicillin-Resistant Staphylococcus aureus Quorum Sensing and Protects Host Skin from Damage. Antimicrob Agents Chemother 64, e00172–20 (2020).

27. Ramsey, M. M., Freire, M. O., Gabrilska, R. A., Rumbaugh, K. P. & Lemon, K. P. Staphylococcus aureus Shifts toward Commensalism in Response to Corynebacterium Species. Front Microbiol 7, 1230 (2016).

28. Caballero, S. et al. Cooperating Commensals Restore Colonization Resistance to Vancomycin-Resistant Enterococcus faecium. Cell Host Microbe 21, 592–602.e4 (2017).

29. Meisel, J. S. et al. Skin Microbiome Surveys Are Strongly Influenced by Experimental Design. J Invest Dermatol 136, 947–956 (2016).

30. Zheng, Q., Bartow-McKenney, C., Meisel, J. S. & Grice, E. A. HmmUFOtu: An HMM and phylogenetic placement based ultra-fast taxonomic assignment and OTU picking tool for microbiome amplicon sequencing studies. Genome Biol 19, 82 (2018).

31. McMurdie, P. J. & Holmes, S. phyloseq: an R package for reproducible interactive analysis and graphics of microbiome census data. PLoS One 8, e61217 (2013).

32. Turner, S., Pryer, K. M., Miao, V. P. & Palmer, J. D. Investigating deep phylogenetic relationships among cyanobacteria and plastids by small subunit rRNA sequence analysis. J Eukaryot Microbiol 46, 327–338 (1999).

33. Wickham, H. ggplot2. (Springer International Publishing, 2016). doi:10.1007/978-3-319-24277-4.

34. Yu, G. Using ggtree to Visualize Data on Tree-Like Structures. Current Protocols in Bioinformatics 69, (2020).

