## Supplemental Figures and Table for "Harnessing diversity and antagonism within the pig skin microbiota to identify novel mediators of colonization resistance to methicillin-resistant *Staphylococcus aureus*"

**SUPPLEMENTAL MATERIAL**

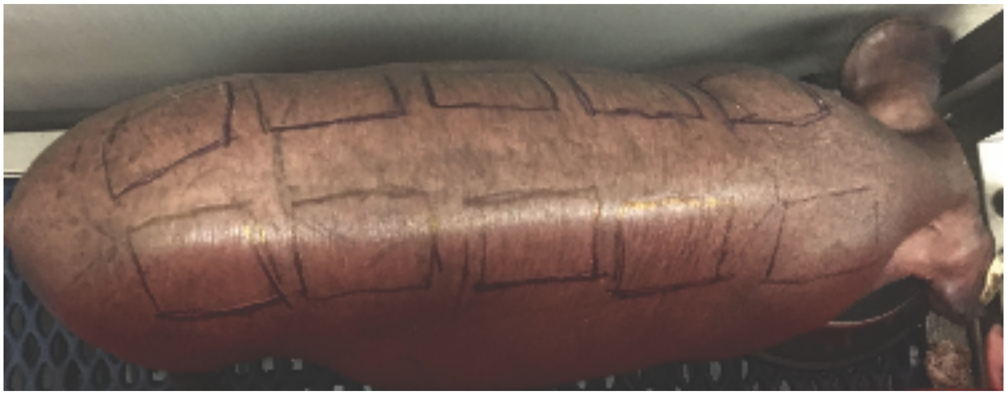

**Figure S1:** Example of pig model for topical treatment and colonization, with 10 patches marked with skin marker.

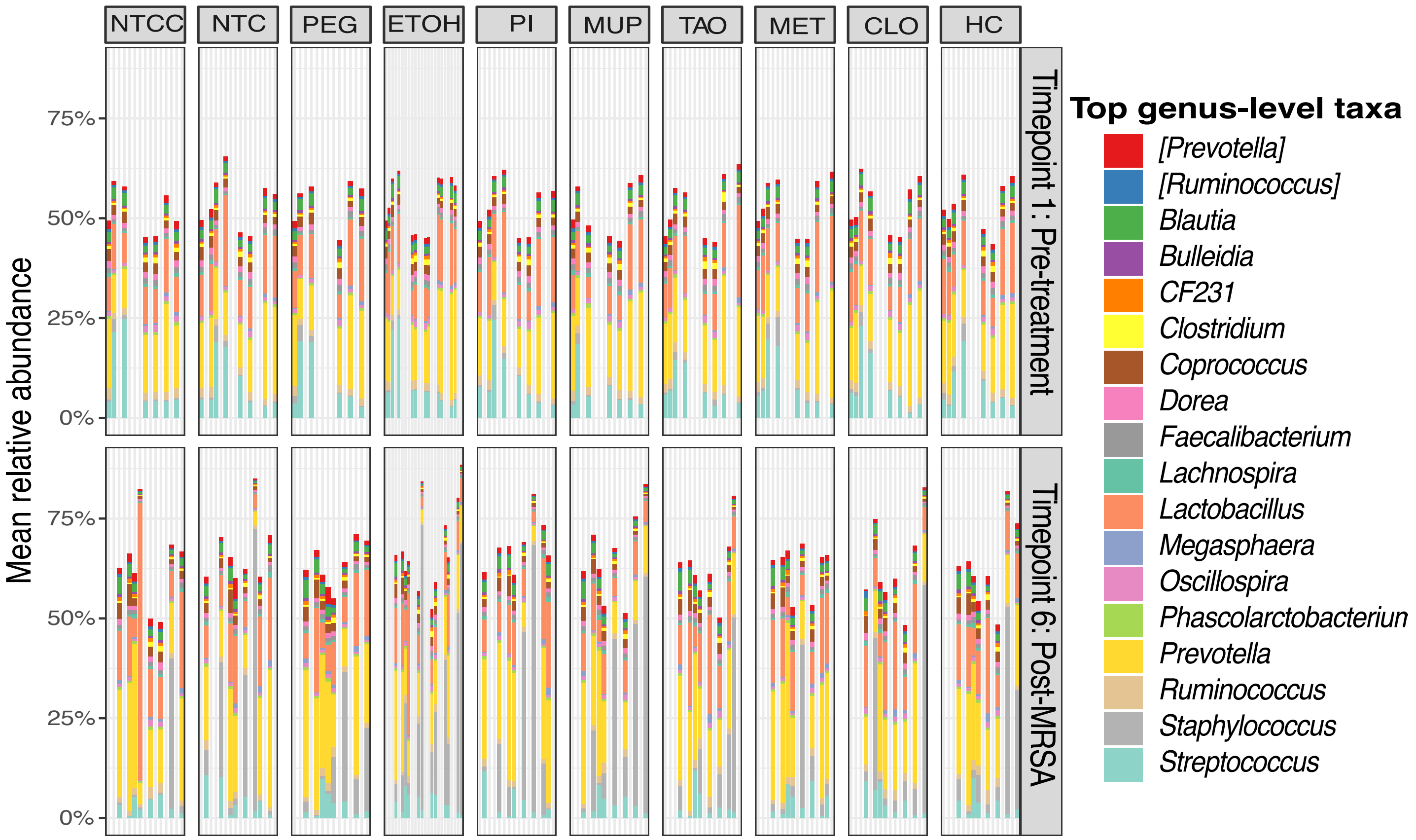

**Supplemental Figure S2:** Individual mean relative abundances (y-axis) of skin bacteria for each pig (x-axis) grouped by treatment. The top panel compares timepoint 1 before any treatments to the bottom panel at timepoint 6 following MRSA colonization.

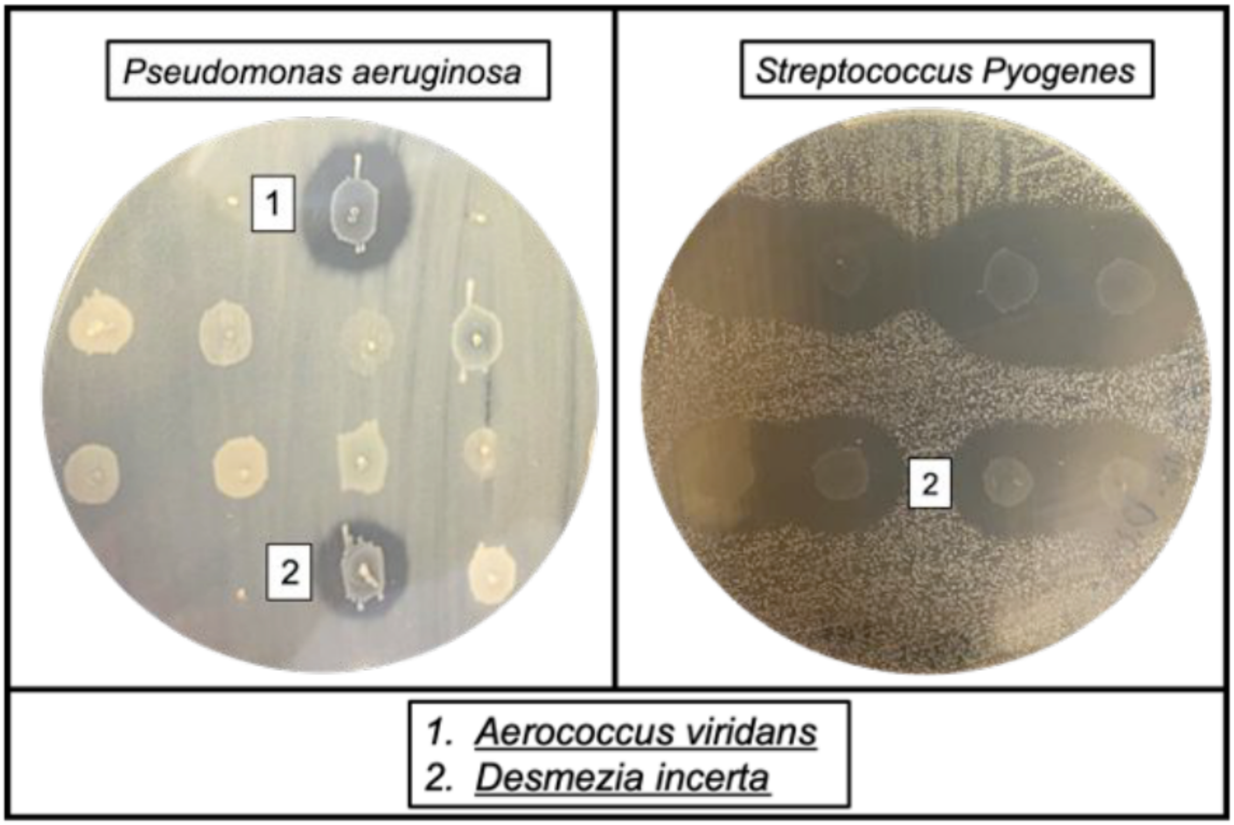

**Supplemental Figure S3:** Additional pathogens inhibited by *A. viridans* and *D. incerta*. A lawn of *Pseudomonas aeruginosa* (left) and *Streptococcus pyogenes* (right) were spotted with pig commensal bacteria. Zones of clearing marked with “1” indicate inhibition by *A. viridans* and “2” indicate *D. incerta*.

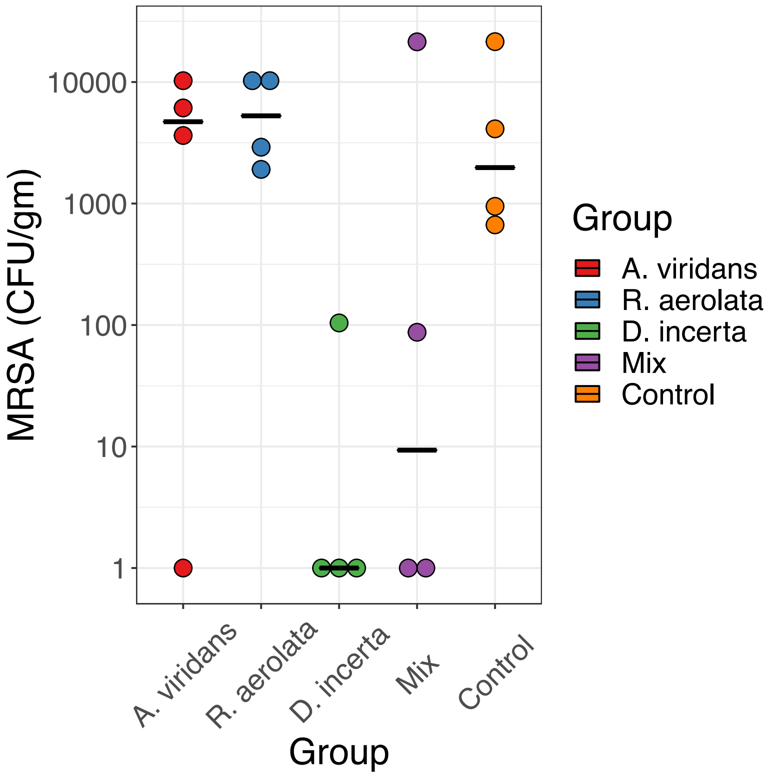

**Supplemental Figure S4:** Pilot experiment of pig isolates and in vivo colonization resistance to MRSA. Inoculum of $4.5x10^7$ CFU of each pig isolate was applied, daily for 2 days. On the $3^{rd}$ day, MRSA inoculum of $1x10^8$ CFU was applied. Twenty-four hours later, skin was collected and CFU MRSA per gram tissue calculated (y-axis) for each treatment group (x-axis).

**Supplemental Table S1: List of unique species isolated from skin of pigs**

| Isolate | Inhibits MRSA? |
|---|---|
| *Acinetobacter lwoffii* | Yes |
| *Aerococcus suis* | |
| *Aerococcus viridans* | Yes |
| *Amycolatopsis alba* | |
| *Bacillus aerius* | Yes |
| *Bacillus altitudinis* | Yes |
| *Bacillus cereus or Bacillus mobiluss* | Yes |
| *Bacillus pumilis* | Yes |
| *Bacillus safensis* | Yes |
| *Bacillus sp or Kocuria carniphila* | Yes |
| *Bacillus subtilis* | Yes |
| *Bacillus tequilensis* | Yes |
| *Bacillus zhangzhouensis* | Yes |
| *Bergeyella porcine* | |
| *Brachyspira sp (muris?)* | |
| *Brevundimonas diminuta* | Yes |
| *Candida guilliermondii* | Yes |
| *Corynebacterium camporealensis* | |
| *Corynebacterium confusum* | |
| *Corynebacterium sp* | |
| *Corynebacterium lipophiloflavum* | |
| *Corynebacterium mycetoides* | |
| *Corynebacterium pollutisoli* | |
| *Corynebacterium glutamicum* | |
| *Corynebacterium sphenisci* | |
| *Corynebacterium spheniscorum* | |
| *Corynebacterium tuscaniense* | |
| *Corynebacterium vitaeruminis* | |
| *Corynebacterium xerosis* | |
| *Cryptococcus magnus* | Yes |
| *Delftia acidovorans* | Yes |
| *Desemzia incerta* | Yes |
| *Enterococcus casseliflavus* | Yes |
| *Enterococcus faecalis* | Yes |
| *Enterococcus saccharolyticus* | |
| *Escherichia vulneris* | Yes |
| *Facklamia hommis* | |
| *Kocuria atrinae* | |
| *Kocuria gwangalliensis* | |
| *Kocuria rhizolphilla* | Yes |
| *Leucobacter chromiiresistens* | |
| *Lysinibacillus fusiformis* | Yes |
| *Microbacterium arborescens* | Yes |
| *Microbacterium liquefaciens* | Yes |
| *Microbacterium maritypicum* | |

| | |
|---|---|
| *Microbacterium oxydans* | |
| *Microbaterium paraoxydans* | Yes |
| *Micrococcus endophyticus* | |
| *Moraxella osloensis* | |
| *Moraxella pluranimalium* | |
| *Neisseria perflavia* | |
| *Paenibacillus sIlvae* | Yes |
| *Pseudomonas azotoformans* | |
| *Pseudomonas chlororaphis* | |
| *Pseudomonas cremoricolorata* | |
| *Pseudomonas fulva* | Yes |
| *Pseudomonas koreensis* | Yes |
| *Pseudomonas oryzihabitans* | |
| *Pseudomonas parafulva* | |
| *Rhodococcus erthropolis* | |
| *Roseomonas gilardii* | |
| *Rothia aerolata* | Yes |
| *Rothia nasisurum* | Yes |
| *Staphylococcus chromogenes* | Yes |
| *Staphylococcus cohnii* | |
| *Staphylococcus divriesel* | |
| *Staphylococcus equorum* | Yes |
| *Staphylococcus gallinarum* | |
| *Staphylococcus haemolyticus* | Yes |
| *Staphylococcus homis* | |
| *Staphylococcus petrasii* | |
| *Staphylococcus sciuri* | |
| *Staphylococcus simulans* | Yes |
| *Staphylococcus sp* | |
| *Staphylococcus sp presumptive xylosus* | Yes |
| *Staphylococcus warneri* | Yes |
| *Stenotrophomonas maltophilia* | Yes |
| *Streptococcus alactolyticus* | |
| *Streptococcus gallolyticus* | |
| *Streptococcus oralis* | |
| *Streptococcus porcorum* | |
| *Streptococcus suis* | |
| *Streptococcus tangierensis* | |
| *Trichosporon asahii* | Yes |
| Isolates were identified by MALDI-TOF Mass spectrometry and 16S rRNA gene sequence | |
| Isolates (37) that inhibited MRSA were preferentially collected. | |
